# Behavioral recovery after a spinal deafferentation injury in monkeys does not correlate with corticospinal sprouting

**DOI:** 10.1101/2020.01.27.921957

**Authors:** Matthew Crowley, Alayna Lilak, Joseph Garner, Corinna Darian-Smith

## Abstract

A long held view in the spinal cord injury field is that corticospinal terminal sprouting is needed for new connections to form that then mediate behavioral recovery. The inference is that more extensive sprouting predicts greater recovery, though there is little evidence to support this. Here we compare behavioral data from two established deafferentation injury models in monkeys, to provide a clear example that sprouting does not track with behavioral recovery.

## Main

Recovery of function following spinal cord injury is notoriously difficult to predict, and experimental models have long used axonal sprouting, and most commonly corticospinal tract (CST) outgrowth, as an anatomical biomarker of functional recovery^1-7^. Here we compared behavioral recovery across two spinal injury models in the macaque monkey, that cut primary afferent inputs exclusively from the thumb, index and middle finger of one hand ^8-11^, to determine the validity of this relationship. Lesions involved either the dorsal rootlets alone (DRL), or in combination with a lesion in the dorsal column (i.e. cuneate fasciculus, DRL/DCL). Both lesions cut a similar (though not identical) afferent population of fibers, and produced a sensory deficit in the opposing digits of the affected hand. Despite the similarities of the two lesions, each is known to induce a dramatically different sprouting response in the spinal cord from the primary somatosensory (S1) and motor (M1) CSTs during the first 6 post-injury months. These data have been published^2,8-12^ (see summary data in **Figures 1D-E**) and set the stage for the current investigation. When the dorsal roots alone are transected, the S1 CST on the lesioned side retracts to 60% of its original terminal territory within the dorsal horn of the cervical cord (**Figure 1D**, green vs orange distributions), and the M1 CST remains robust and largely unchanged from its normal range (**Figure 1E**, green vs orange). In contrast, following a DRL/DCL, both the S1 and M1 CST terminals sprout dramatically (and bilaterally) beyond their normal range within the cervical and thoracic cord (**Figure 1D-E**, compare blue with green territories)^2,8,9,11,12^. The injury models used were easily replicated, and ideal for testing the relationship between terminal sprouting and functional recovery extent^1,13^. Injuries that transect the CSTs directly (e.g. hemisections, or contusions) are clinically relevant, but also more difficult to precisely reproduce, which makes it challenging to interpret the impact of terminal sprouting on behavioral recovery.

**Figure 1.**
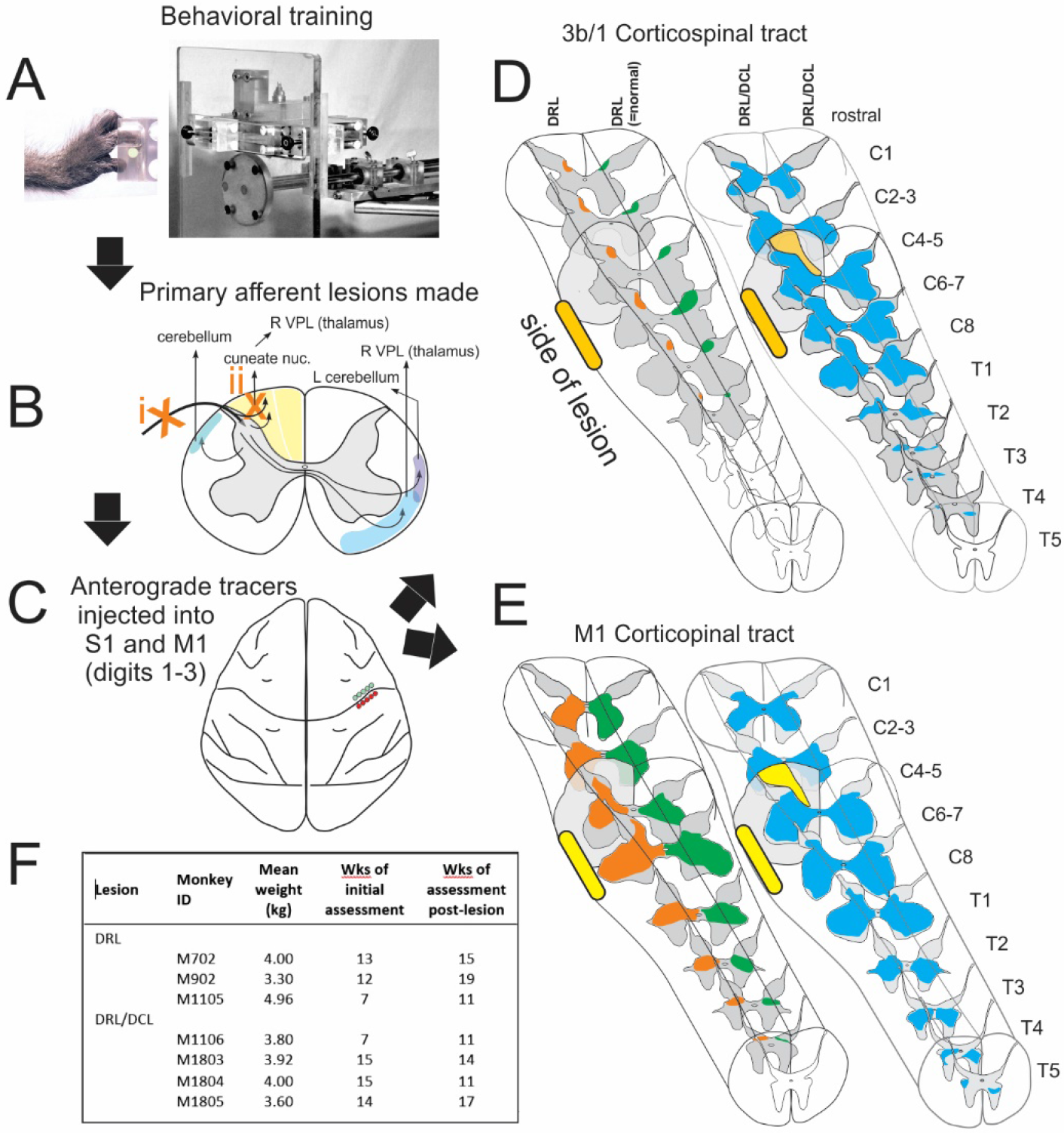
**A-C** Summarizes the experimental design of the study, with the behavioral manipulandum/apparatus used to assess reach-grasp-retrieval (**A**), the location of, and pathways impacted by the two lesion models (**B**), and the placement of tracers within the sensorimotor cortical regions during a craniotomy (**C**), which labeled corticospinal projections to the cervical and thoracic cord. These anatomical findings have been published ^2,8,9,11^ and they provide the context for this report and behavioral analysis. (**D-E**) Summarizes these published data, and clearly show the dramatic differences in terminal sprouting responses in the two lesion groups^2,8,9,11^. (**F**) provides specific details for the 7 monkeys used.

**Figure 2.**
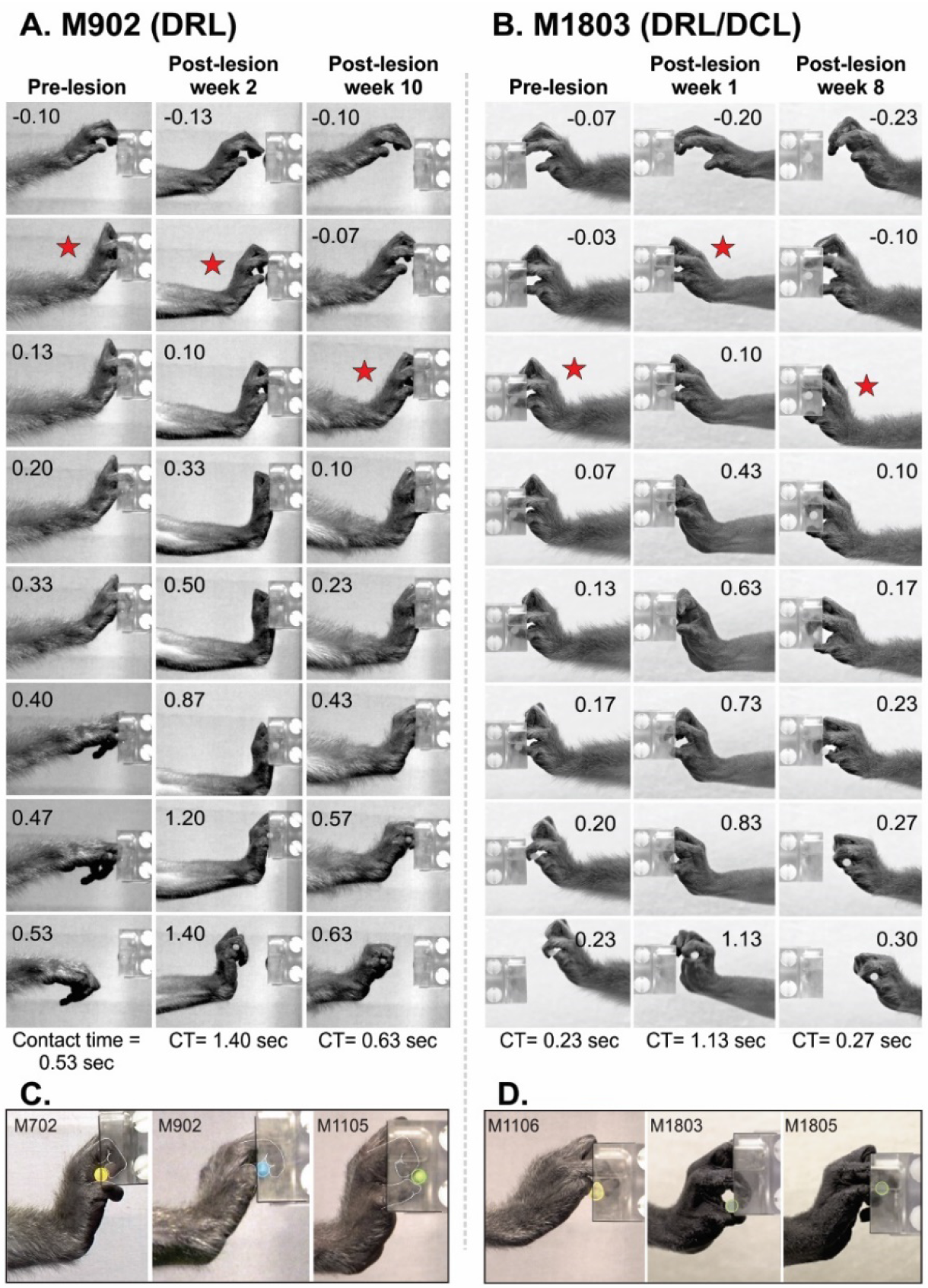
Representative manual retrieval stratagems used during behavioral assessment. (**A-B**) Three different frame sequences each from monkeys M902 (DRL) and M1803 (DRL/DCL), showing the predominant stratagem used prelesion (column 1) and postlesion (columns 2 and 3). Contact times for each sequence are given at the bottom of each column. The times for each frame are indicated, with red stars illustrating the initial digit contact with the object and the point at which contact times begin. Prior to the lesion (column 1), performance was smooth and efficient. The target was contacted simultaneously with the opposing distal pads of the thumb and index finger, and the object was retrieved from the clamp. In column 2 (2 weeks postlesion for M902 and 1 week postlesion for M1803), performance was visibly impaired, and retrieval took much longer than it had prior to the lesion. Retrieval stratagem was also different, as initial contact with the object no longer involved simultaneous opposition of the thumb and index finger. Instead, only the distal pad of the index finger made initial contact, which then scooped the object until it contacted the distal pad of the thumb. Several attempts were needed to dislodge the object from the clamp, after which the object was supported by the dorsum of the distal segment of the thumb and the palmar pads of the curled index finger. Placement of both digits were abnormal. There was also a profound extension of the wrist, which was not seen prelesion. In column 3 (10 weeks postlesion for M902 and 8 weeks postlesion for M1803), a similar scooping motion was used, though the overall proficiency of the performance had increased and the contact times decreased. Although a compensatory stratagem was adopted to retrieve the object, this new strategy produced the same efficiency seen prior to the lesion. (**C-D**) Alternate stratagems used post-lesion. Monkeys adopted numerous compensatory stratagems, as illustrated in order to successfully grasp and retrieve the object from the clamp.

**Figure 3.**
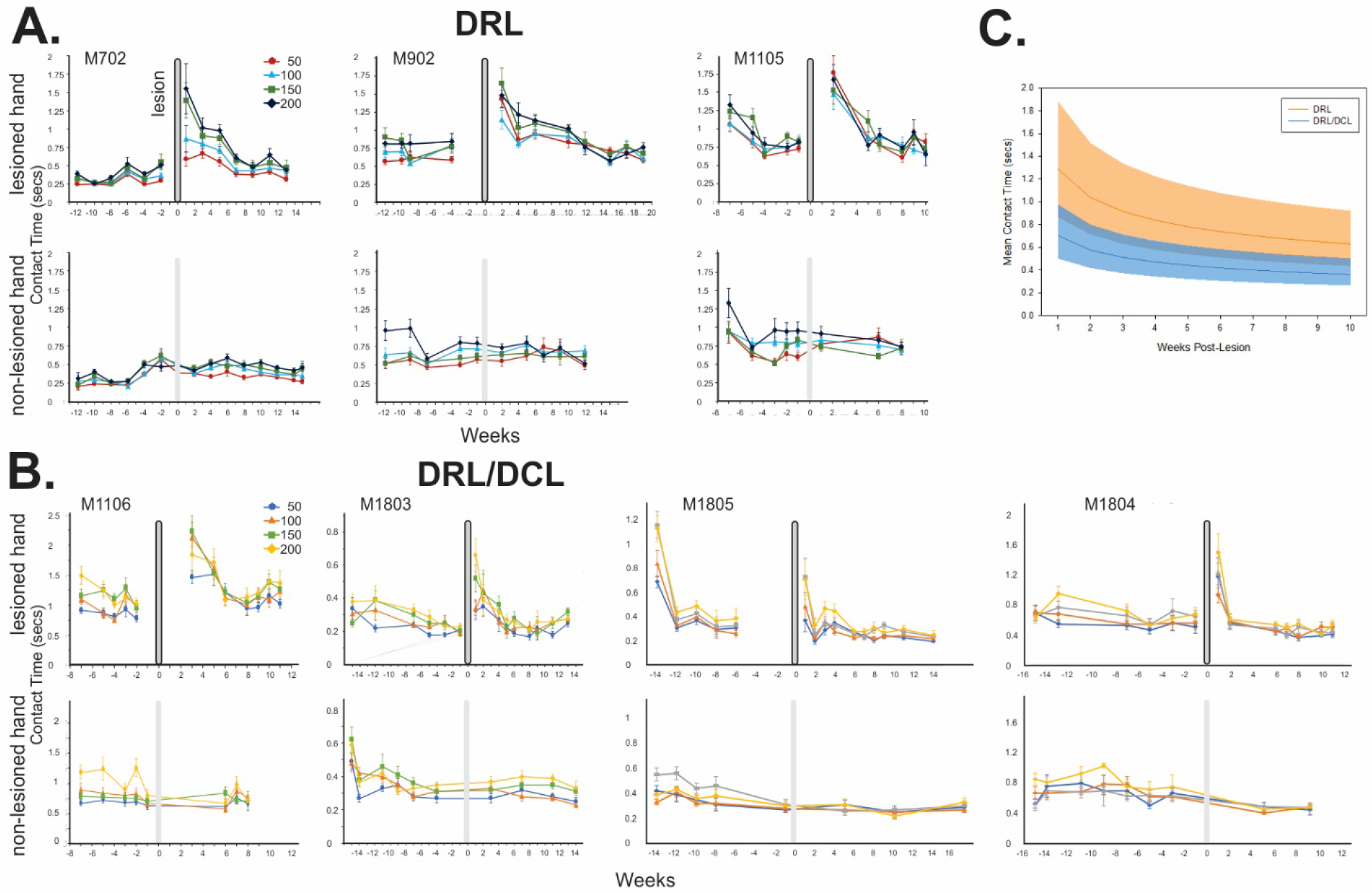
(**A-B**) Contact time data showing the raw data for lesioned and non-lesioned hands in monkeys from each lesion group, and (**C**) a summary of data from the lesioned hand in all monkeys. Recovery of function is shown by a decrease in contact time following the lesion. The graph shows the line of best fit for the two lesion types, figured for the average force. The shaded area is the +/- SE of each line. Importantly, the lines would have to be separated by approximately 3 standard errors to be significantly different. The decrease in contact time is highly significant, but the difference between the two curves is not significant. The analysis was performed in log-log space to model these curves plotted in linear space on the figure.

Given our findings, and the different CST responses following the two lesions, we hypothesized that there is no clear link between the CST terminal sprouting extent and the post-injury behavioral recovery.

All monkeys in the current study underwent the same experimental sequence shown in **Figure 1**, and published elsewhere^8,9,11,12^. Briefly, all were trained and assessed in a reach-grasp-retrieval task requiring sensory feedback, and once performance had stabilized, a laminectomy was performed to make the lesion. Electrophysiological recordings were made in dorsal rootlets in C5-T1 to identify and target only the rootlets with detectable cutaneous RFs on the thumb, index and middle fingers (D1-D3). This is necessary for lesion replication, due to substantial inter-animal variability of inputs (segmentally)^14^. In monkeys receiving a DRL/DCL, the DCL was always placed at the rostral border of the DRL, and only the cuneate fasciculus was cut, as described elsewhere^8,9,11^. Once lesioned, behavioral data were collected (3-5 days per week) for a minimum of three additional months. A craniotomy was then made contralateral to the lesion, to record and locate the reorganized D1-D3 region of the primary somatosensory cortex, so that tracers could be injected to label corticospinal projections to the cord^8,9,11^. Monkeys were euthanized 6-7 weeks after the craniotomy.

Behavioral data were obtained for both the lesioned and non-lesioned hands throughout the assessment period (**Figure 3**). This showed interhand, and interanimal differences in performance times, as well as stable and consistent performance metrics in the non-lesioned hand in all monkeys throughout.

The reach-grasp-retrieval task has been described elsewhere in detail ^15^. Briefly, monkeys reached to retrieve a pellet from one of four clamps (on a rotating turret) at a set distance in front of them. Each clamp held an identical treat at a different resistive force (0.5, 1.0, 1.5 and 2.0 Newtons), which made trials dependent on sensory feedback for effective retrieval. Monkeys could not see the clamp after the initial reach.

Baseline performance was taken as the data obtained during the last two weeks of pre-lesion training, since monkeys typically followed a learning curve during the early training period before their performance stabilized. Two parameters were used to assess the deficit and recovery of function in the hand: (1) contact time (CT), or the time from initial contact with the pellet to its removal from the clamp, and (2) the digit stratagem used to grasp and successfully retrieve the pellet. Representative examples of the behavioral consequences of each lesion are shown in **Figure 2**. All monkeys showed an initial deficit, and a subsequent recovery of digit function during the early post-lesion weeks. Monkeys performed the retrieval task pre-lesion using a precision grip involving the opposition of the distal pads of digits 1 and 2. After either lesion, however, performance was clumsy. Monkeys could preshape the affected hand, but were unable to accurately locate the target using the opposing distal digit pads once they reached the clamp, or to apply the correct force needed to effectively retrieve the target. This meant that contact times were initially significantly longer than baseline (**Figure 3**). Over 5-8 weeks, however, all animals developed novel and effective grip strategies (**Figure 3**), which allowed them to perform the task at pre-lesion speed. This was functional compensation, rather than a complete restoration of the original strategy. All DRL and DRL/DCL animals (targeting D1-D3), without exception, have followed a similar deficit and post-injury recovery of hand/digit function.

For the statistical analysis, we used data from all 7 monkeys and compared CTs for the two lesion types (DRL versus DRL/DCL), over pre-lesion and post-lesion weeks, for the different clamp forces (0.5, 1.0, 1.5, and 2.0 Newtons). Only data from the side of the lesion was assessed statistically, as contact times, as well as stratagems were unaffected in the contralateral hand throughout the assessment period (**Figure 3**).

Contact time was log-normal distributed (as is typical for latencies), so CTs were logged and averaged for analysis. To test whether lesion type affected recovery post lesion, we performed a repeated measures REML mixed model in JMP 14 Pro. Monkeys were nested within lesion type, and treated as a random effect. Force and week were treated as continuous variables. Week was logged to model curves of decreasing return. Interactions between lesion type, and force and lesion type and week were tested to determine whether the effect of force or week differed between the lesions. Appropriate error terms for repeated measures mixed models were included.

We found that the change in contact times (i.e. slope) did not differ significantly between the two lesion types (F_1,5.00_=1.322; P=0.3022; **Figure 3**). CT increased significantly with force (F_1,157_ = 33.05; P<0.0001), but this effect did not differ between lesion types (F_1,157_=0.0557; P=0.8138). CTs also decreased significantly with week post lesion (F_1,157.1_ = 182.7; P<0.0001), and this effect did not differ between lesion types (F_1,157.1_=0.2327; P=0.6302; **Figure 3**).

Given that the two lesions differed so dramatically with respect to S1 and M1 corticospinal tract responses, but not with respect to the monkey’s hand function, our findings support our hypothesis and provide the first clear demonstration that the extent of CST terminal sprouting does not predict behavioral recovery. We demonstrate that more sprouting does not predicate better behavioral outcome. This means that CST sprouting is not in itself a good biomarker of behavioral recovery. Our findings do not suggest that axon terminal growth is not important (or even critical) in the formation of new connections. Clearly some outgrowth is needed for the formation of new synapses, but this may only require very localized sprouting. The relationship is clearly complex and more nuanced, involving all levels of the neuraxis, the mix of peripheral to central pathways, and factors such as the post-lesion timepoint at which responses are assessed, synaptic connectivity changes, the inflammatory response^16^, and molecular and cellular changes^13^. Excess axonal growth cannot be used to infer a better behavioral outcome, and future studies are needed to establish more reliable predictors of behavioral recovery.

## Online Methods

### Subjects

Subjects were 7 young adult (3-4 years old with an average weight of 3.9 ± 0.5kg) colony bred (Charles River) male monkeys (*Macaca cynomolgus*). Monkeys were housed individually at the Stanford Research Animal Facility, in four unit cages (64×60×77cm, depth x width x height per each unit) in a room with other monkeys. They were kept to a 12 hour light/dark cycle and the room was maintained at 72-74°F. ARRIVE guidelines were followed, with the exception that only male monkeys were used. This was due to availability and a lack of gender differences with respect to hand function, sensorimotor pathways and recovery following SCI.

Animal procedures were carried out in accordance with National Institutes of Health guidelines and the Stanford University Institutional Animal Care and Use Committee.

### Behavioral assessment

All monkeys were matched for species, age, and sex, and all had identical DRLs or combined DRL/DCLs that only affected the thumb, index and middle fingers (D1-D3) in one hand (see **Figure 1**). All lesions were unilateral, and made on the side of the dominant hand. Hand preference was determined over 2-3 weeks prior to the laminectomy. Monkeys were handed fruit or other small food items, and the preferred hand for reaching and grasping the food was scored over dozens of reaches during this time period. All monkeys had identical behavioral training routines. As such, the monkeys in each lesion group were otherwise indistinguishable.

The reach-grasp-retrieval task involved a natural movement, so that monkeys learned quickly and performed at a consistent speed within 4-6 weeks. Monkeys were trained to sit in a plexiglass box and reach through a window (located on either the right or left depending on the hand being tested), to retrieve a candy pellet held in one of 4 clamps. Each clamp held an identical pellet at one of 4 different forces (0.5, 1.0, 1.5 or 2.0 Newtons), and clamps (which were visually indistinguishable to the monkey), were presented pseudo-randomly. Training sessions were filmed using a high shutter speed (30 fps at 1/500s) digital camcorder (Canon XF200) and data analyzed offline using Edius Pro 9 (Grass Valley) software. ‘Contact time’ was the time interval between first contact with the target pellet and its successful displacement from the clamp.

Data were obtained from a minimum of 20 trials per week, pooled from 3-5 training sessions of 20 minutes, and conducted at the same time each day. Monkeys had chow and water available ad libitum, but fruit was provided after training and candy pellets were only available during training.

### Surgical Procedures

All monkeys underwent two surgical procedures, including a laminectomy to make the lesion, and a craniotomy to place anterograde tracer injections into reorganized primary sensory and motor cortex. The latter labeled CST terminals within the spinal cord, and the anatomical data have been described and published elsewhere^8,9,11^.

All surgical procedures involved initial sedation with ketamine hydrochloride (10mg/kg), and maintained under gaseous anesthesia (isofluorane, 1-2% / O_2_). Atropine sulfate (0.05mg/kg), buprenorphine (0.015mg/kg) and the antibiotic cefazolin (20mg/kg) were also given, and Normasol-R was infused intravenously throughout surgery to maintain fluid balance. Dexamethasone (0.25mg/kg) was also given before craniotomy procedures to minimize brain edema.

Physiological signs (i.e. blood pressure, heart rate, pulse oximetry, capnography, and core temperature) were continuously monitored throughout surgery to ensure a proper depth of anesthesia. A post-operative analgesic (buprenorphine, 0.015mg/kg) was given after surgery and monkeys were typically awake and alert within one hour. Oral meloxicam (0.1mg/kg) was administered for 3-5 days post-surgery, and Buprenorphine (0.05mg/kg, oral route) was also given as indicated in the days following surgery.

## Acknowledgements

This work was supported by the National Institute of Neurological Disorders and Stroke (R01 NS091031 to CD-S). We would like to thank Karen M. Fisher for her help with surgical procedures, manuscript comments, and neuroanatomical analyses, and Cholawat Pacharinsak, and Benjamin Franco for their help with anesthesia.

## Data Availability Statement

The data that support the findings of this study are available from the corresponding author upon reasonable request.

